# In vitro toxicity of LiTFSI on Human Renal and Hepatoma Cells

**DOI:** 10.1101/2023.08.15.553404

**Authors:** Xing Zhang, Mia Sands, Mindy Lin, Jennifer Guelfo, Joseph Irudayaraj

## Abstract

We evaluate the cytotoxicity, intracellular redox conditions, apoptosis, and methylation of DNMTs/TETs upon exposure to LiTFSI, a novel PFAS compound commonly found in lithium-ion batteries, on human renal carcinoma cells (A498) and hepatoma cells (HepG2). The MTT assay showed both PFOS and LiTFSI had a dose-dependent effect on A498 and HepG2, with LiTFSI being less toxic. Intracellular redox conditions were assessed with a microplate reader and confocal, which showed a significant decrease in ROS levels and an increase in SOD content in both cells. Exposure to LiTFSI enhanced cell apoptosis, with HepG2 being more susceptible than A498. Quantitative analysis of mRNA expression levels of 19 genes associated with kidney injury, methylation, lipid metabolism and transportation was performed. LiTFSI exposure impacted kidney function by downregulating Acta2 and upregulating Tgfb1, Bcl2l1, Harvcr1, Nfe2l2, and Hes1 expression. LiTFSI exposure also affected the abundance of transcripts associated with DNA methylation by the expression of TET and DNMT genes. Furthermore, LiTFSI exposure induced an increase in lipid anabolism and alterations in lipid catabolism in HepG2. Our results provide new insight on the potential role of a new contaminant, LiTFSI in the regulation of oxidative stress, apoptosis and methylation in human renal carcinoma and hepatoma cells.

## Introduction

Per and Polyfluoroalkyl Substances (PFAS) are fluorinated substances that contain at least one fully fluorinated methyl or methylene carbon atom (Evich et al., 2022). PFAS usually contains a hydrophobic and oleophobic carbon chain structure and a hydrophilic functional group (Zhang et al., 2022). The structure of PFAS encompasses C-F bonds which has a high bond energy and strong polarity (Huang and Jaffé, 2019). The amphiphilicity and C-F bonds contribute to the stability, surface activity, hydrophobic, and oleophobic properties of PFAS. PFAS have been widely used in the production of various industrial and household products, such as the coatings on non-stick pans, hydrophobic and oleophobic coatings, foam fire extinguishing agents, and surfactants (Kucharzyk et al., 2017). Due to the highly stable C-F bonds, PFAS are recalcitrant, resulting in their lasting persistence in the environment and the human body (Fauconier et al., 2020).

The widespread use of PFAS has raised concerns on the toxic effects and human health risks. Numerous studies have shown that PFAS are widely prevalent in the environmental media and organisms including air (Chen et al., 2019), sludge (Higgins et al., 2005), surface water (Habibullah-Al-Mamun et al., 2016), soil (Ahrens et al., 2011), and even polar ice sheets (Kannan et al., 2001), wild animals (Hong et al., 2015), and humans (Yeung et al., 2006). After entering the human body, PFAS chemicals accumulate in organs via the blood circulatory system, causing damage and endocrine disruption (Ahmad et al., 2021; Forsthuber et al., 2020).

Our specific focus is the in vitro toxicity of LiTFSI (CAS No. 90076-65-6), also known as lithium bis(trifluoromethanesulfonyl)imide, LiNH(CF3SO2)2, a lithium salt with a weak coordination anion (Kubota et al., 2010). Due to its superior electrochemical stability and conductivity, LiTFSI is a popular choice as a composite polymer electrolyte material, often utilized in lithium-ion batteries and organic electrolyte lithium salts (Zhang, 2022). LiTFSI is considered to be a type of PFAS due to its similarity in molecular structure (Gilbert et al., 2017). It contains a perfluoroalkyl chain and shares many parallels with other PFAS and is also highly stable and persistent in the environment. Despite its prolonged and widespread use in the energy sectors, the toxic effects of LiTFSI have yet to be studied; the potential adverse effects on human health and the environment are largely unknown. It is believed that it may share some of the same properties and potential risks as other PFAS. Our work investigates the potential toxic effects of LiTFSI in relation to the kidney and liver, in vitro.

Livers and kidneys are some of the most common organs for contaminant accumulation. PFAS exposure induced hepatomegaly (Wen et al., 2020), chronic kidney disease (CKD) (Li et al., 2022) and cancer in mice. PFAS is also known to induce oxidative stress, which can lead to cell death and tissue damage in severe cases (Lu et al., 2019; Ojo et al., 2021; Wielsøe et al., 2015). Exposure to certain PFAS has been associated with liver damage (Costello et al., 2022), including inflammation, hepatocyte necrosis, and liver tumors. Additionally, PFAS exposure has been shown to decrease liver weight, alter liver enzymes and lipid metabolism, and induce oxidative stress (Roth et al., 2021). A study of individuals exposed to PFAS through contaminated drinking water found a positive association between PFAS exposure and increased liver enzymes (Sen et al., 2022). Exposure to PFAS has been linked to decreased kidney function, such as decreased glomerular filtration rate (GFR) (Verner et al., 2015)and increased albuminuria (Jain and Ducatman, 2022), which are signs of kidney disease.

We will investigate the short-term toxicity of LiTFSI on cell viability, reactive oxygen species (ROS) and superoxide dismutase (SOD) levels, apoptosis, cell cycle, and gene expression related to methylation. We expect our foundational work to provide the basis for future research on the use and mitigation of PFAS compounds such as LiTFSI.

## 2 Materials and Methods

### 2.1. Chemicals and test reagents

The lithium salt form of LiTFSI, Lithium bis(trifluoromethanesulfonyl)imide (CAS No. 90076-65-6) 99.99% grade was purchased from Sigma-Aldrich Corporation (St. Louis, MO, USA). Perfluorooctane sulfonate (PFOS) was purchased from SynQuest Labs (Alachua, FL, USA). LiTFSI and PFOS were dissolved in dimethyl sulfoxide (DMSO; CAS No. D8418) from Sigma-Aldrich to obtain a stock solution of 0.1□M.

### 2.2 Cell culture

The HepG2 (ATCC HB-8065) and A498 (ATCC HTB-44) cell lines were obtained from the Cancer Center at Illinois, University of Illinois Urbana-Champaign. HepG2 cells were cultured in Dulbecco’s Modified Eagle’s Medium (ATCC 30–2002; Manassas, VA, USA) supplemented with 10% fetal bovine serum (FBS, (Gibco™ 10082147; Thermo Fisher Scientific; Waltham, MA, USA) and 1% Penicillin Streptomycin Solution (REF 30–002-Cl; Corning, NY, USA). A498 was cultured in Eagle’s Minimum Essential Medium (ATCC 30–2003; Manassas, VA, USA) supplemented with 10% FBS and 1% Penicillin Streptomycin Solution. Culture conditions for all cells were at 37 °C in a humidified 5% CO2 atmosphere. All experiments were performed independently in triplicate.

### 2.3 MTT assay

Cell growth and proliferation assay at 0, 50, 100, 150, 200 or 250 μM of LiTFSI and PFOS were measured with the MTT assay kit (Invitrogen, Cat. No. M6494). Cells were plated into a 96-well microplate (Corning Incorporated, New York, USA) and incubated at 37 °C in 5% CO2. Each data point represents measurement from three replicate wells. The cells were cultured to 30% to 40% confluence before corresponding treatments and cultured for another 24 h or 48 h to reach a final confluence of 50% to 90%. After exposure to chemicals, 10 μl of MTT reagent was added to each well with 90 μl of serum-free medium and incubated for another 4 h at 37 °C. MTT reagent was then added to the medium to stop the reaction and the absorbance, at 570 nm, was measured using a microplate reader (Synergy HT, BioTek; Winooski, VT, USA).

### 2.4 ROS/Superoxide detection

The production of ROS and superoxide were measured in A498, and HepG2 cells by ROS/Superoxide Detection Assay Kit (Cell-based) (ab139476; Abcam; Waltham, MA, USA). Cells were loaded with the ROS/Superoxide Detection Mix at 37°C for 30-60 min in the dark. For confocal measurements the detection mixture was removed from glass slides and cells were washed gently with wash buffer and observed under confocal microscopy with fluorescein (Ex/Em = 490/525nm). For microplate reading, cells were washed and incubated with chemicals in a 96-well plate for 4 h and fluorescence was measured with standard fluorescein (Ex/Em=488nm/520nm) and rhodamine (Ex/Em=550nm/610nm) filter sets at endpoint mode.

### 2.5 Apoptosis assay

To investigate the effects of LiTFSI and PFOS on apoptosis, A498 and HepG2 cells were cultured with chemicals in 6-well plates. After 24 h, plates were harvested and incubated. A TACS® 2 TdT-Fluor *In Situ* Apoptosis Detection Kit (4812-30-K, Minneapolis, MN, USA) was used per manufacturer’s instruction to determine the level of cell apoptosis. Apoptotic cells were analyzed using a BD LSR Fortessa CMtO Analyzer equipped with HTS (High Throughput Sampler) flow cytometer.

### 2.6 Cell cycle assay

Cells in the logarithmic growth phase were incubated with serum-free medium for 6 h to arrest all cells at G0/G1 phase, followed by exposure to different concentrations of LiTFSI. After 48 h, the cells were fixed with pre-cooled absolute ethanol, resuspended in RNaseA, and stained with PI. Cell cycle was analyzed using a BD LSR Fortessa CMtO Analyzer equipped with an HTS flow cytometer.

### 2.7 Gene expression analysis

Total RNA was isolated using GeneJET RNA Purification Kit (K0731, Thermo Fisher Scientific; Waltham, MA, USA) and DNA-free™ DNA Removal Kit (AM1906, Thermo Fisher Scientific; Waltham, MA, USA) was used to remove the genomic DNA. The quantity and quality of extracted RNA were tested with NanoDrop One. cDNA synthesis was performed by high-capacity cDNA Reverse Transcription Kit (Applied Biosystems REF 4368814; Waltham, MA, USA). Analysis of mRNA was performed by qRT-PCR using SYBR Green PCR Master Mix reagents (Applied biosystems REF 4367659; Waltham, MA, USA) according to manufacturer’s specifications, with a Step One Plus RealTimes PCR System (Applied Biosystems, Waltham, MA, USA). Relative quantification was obtained by normalization to GAPDH expression levels. Reactions were run in triplicate. The expression levels of mRNAs were determined by the 2-ΔΔCt method for relative quantification of gene expression. All primer sequences were listed in Table S1.

### 2.8 Statistical Analysis

The data obtained was analyzed using SPSS software, while the graphical illustrations were generated using GraphPad Prism 9 Software. The data was expressed as the mean ± standard error of the mean (SEM), and the statistical significance of the difference between the two experimental groups was determined using Student’s t-test. A significance level of P < 0.05 was considered statistically significant.

## 3. Results

### 3.1 Cytotoxicity assessment

The cytotoxic effect of LiTFSI on human carcinoma renal cell lines, A498 and further on human hepatoma cells HepG2 were first assayed using the MTT reagent. PFOS was used as a positive control. Based on a previous study with PFOS at 300□μM (resulting in 50% cell death) and 24 h of exposure (Florentin et al., 2011), we chose a concentration of up to 250□μM to maintain cell viability after 48□h of treatment. Two cell lines were incubated with different concentrations (50, 100, 150, 200 or 250 μM) of the chemicals for 24 h and 48 h.

Cell viability decreased upon treatment with LiTFSI and PFOS in a dose-dependent manner (Figure 2). In A498 cells, the concentration of the two chemicals at 100 μM had no effect on cell viability. The 24 h and 48 h cell viability of LiTFSI treatment at 150 μM decreased to 88% and 82%, while the PFOS treatment was 72% and 67% respectively. HepG2 cells were significantly more sensitive to LiTFSI and positive control. HepG2 cells exposed to PFOS exhibited 60% and 51% more cell death than LiTFSI at 150 μM. The viability of two cells exposed to DMSO was not significantly different from the untreated control group, hence only the blank was used as the negative control.

**Figure 1.**
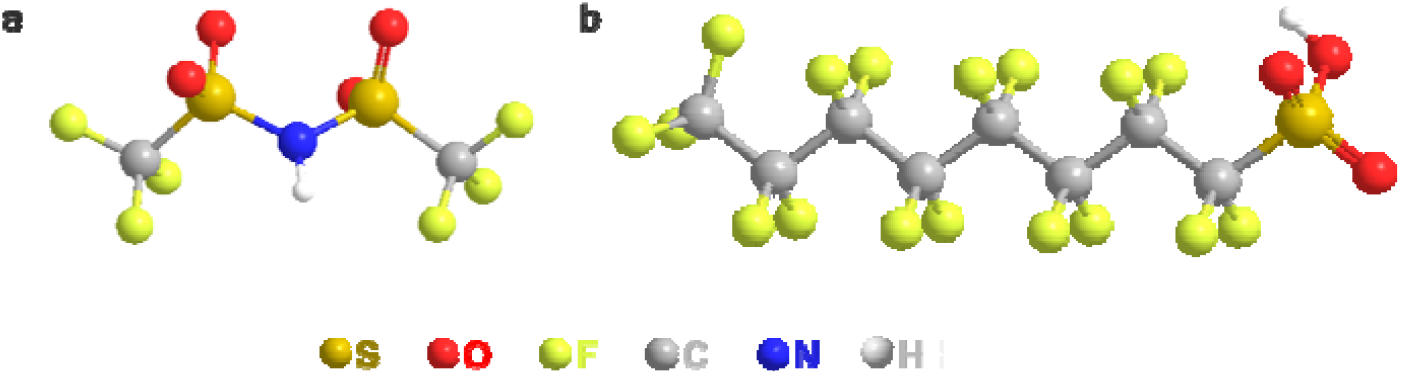
Chemical structure of (a) LiTFSI and (b) PFOS.

**Figure 2.**
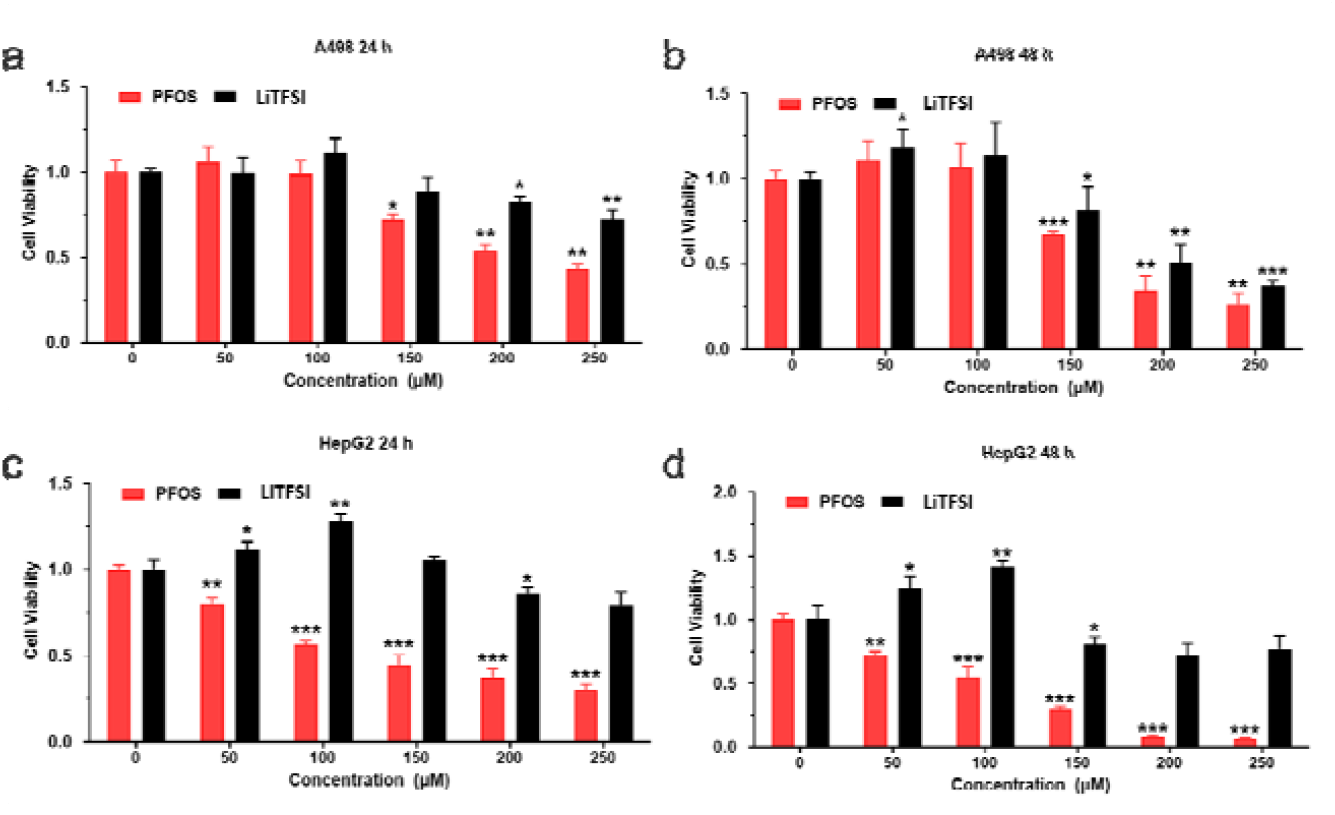
Cytotoxicity assessment of PFOS and LiTFSI in A498 and HepG2 cells. Evaluation of viability of A498 exposed to PFOS and LiTFSI (a) 24 h and (b) 48 h by MTT assay. Viability of HepG2 exposed to PFOS and LiTFSI (c) 24 h and (d) 48 h by MTT assay. Experiments were repeated three times. *p < 0.05 compared with untreated controls(NC).

The concentration of DMSO used in all treatments and controls was kept below 0.4% v/v to minimize its potential influence on cell response. Figure S1 demonstrated that a concentration of 1.6% v/v DMSO had negligible effects when evaluated with MTT assay.

Our data collectively suggest that LiTFSI and PFOS have cytotoxicity on human carcinoma renal cells and hepatoma cells. LiTFSI was less toxic and had higher cell survival after exposure. HepG2 cells were more sensitive to both chemicals.

### 3.2 Intracellular redox conditions

The redox effects of LiTFSI on A498 and HepG2 were assayed using the microplate (Figure 3a) and confocal imaging (Figure 3b). After exposure to A498 and HepG2 cells, the stimulation of ROS and SOD production by LiTFS was measured.

**Figure 3.**
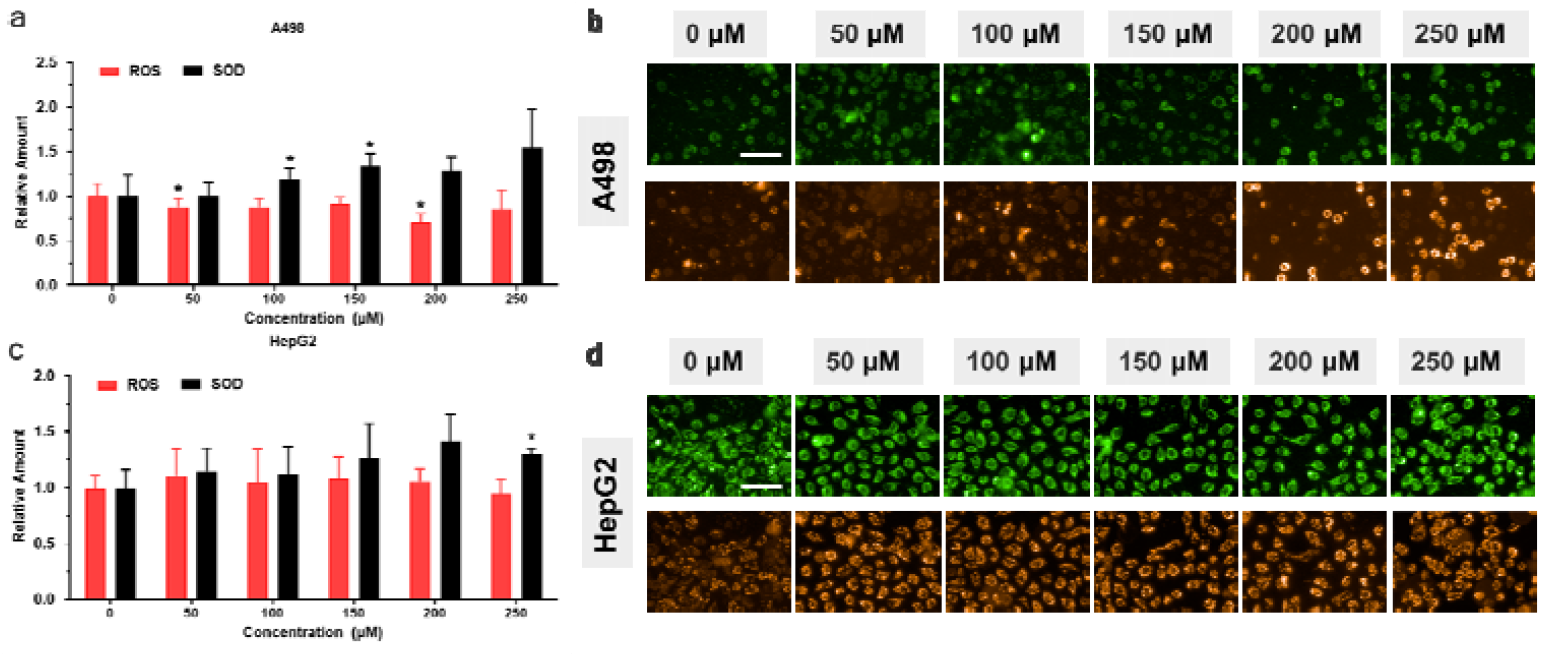
Reactive oxygen species generation and cytosolic superoxide dismutase (SOD) activity of LiTFSI in A498 and HepG2 cells. Experiments were repeated three times. *p < 0.05 compared with untreated controls (NC).

The results of the microplate assay showed a marked reduction in ROS levels in A498 cells. Upon exposure to a concentration of 200 μM of LiTFSI, a significant decrease in ROS levels (by 29%) was noted. Additionally, a simultaneous increase (54%) in SOD content was observed at a concentration of 250 μM, demonstrating a dose-dependent effect. Confocal microscopy results were consistent with the microplate assay, demonstrating an enhancement of red fluorescence, representing an increase in SOD content.

The results of HepG2 revealed a phenomenon that was similar to that observed in A498 cells. At a concentration of 250 μM, a decrease in ROS levels by up to 6% was noted while a corresponding increase in SOD content was 29% compared to the control group. These findings provide insight into the potential role of LiTFSI in regulating cellular oxidative stress in the two cell lines. Further investigation and analysis are necessary to fully understand the underlying mechanisms and implications of these findings.

### 3.3 Cell apoptosis

Cell apoptosis was evaluated by flow cytometry after exposure to LiTFSI for 24 h. Results showed a remarkable increase in the percentage of apoptotic cells in the two cells (p < 0.05, Figure 4a, Figure 4c and Figure S2). HepG2 cells exhibited a higher susceptibility to apoptosis compared to A498 cells. At a concentration of 250 μM, the apoptosis rate of A498 cells was 8.1%, representing a 146% increase compared to the control group. Meanwhile, the apoptosis rate of HepG2 cells was 17.02% under the same conditions.

**Figure 4.**
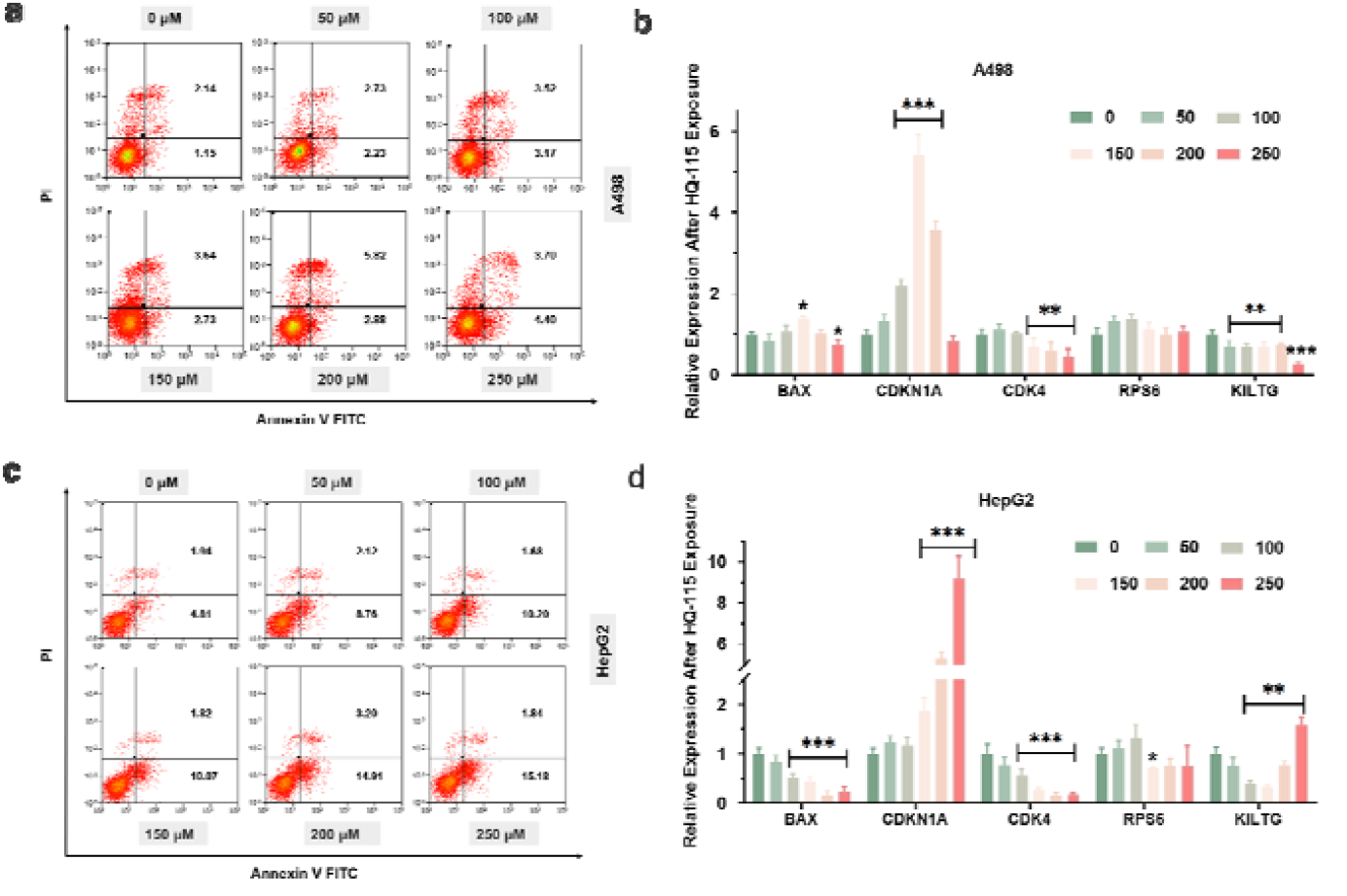
Cell apoptosis of A498 and HepG2 upon exposure to LiTFSI. (A) A498 (B) HepG2 apoptosis distributions detected by flow cytometry. Cells were exposed to LiTFSI for 24 h. (C) A498 (D) Relative expression of HepG2 cell apoptosis and proliferation related genes after 48 h of exposure to LiTFSI. Experiments were repeated three times. *p < 0.05 compared with untreated controls(NC).

Due to the influence of LiTFSI on cell viability and apoptosis, several genes related to apoptosis and proliferation were evaluated. Results showed that the expression of *BCL2*-associated X (*BAX)* in A498 cells initially increased and then decreased, reaching the highest expression at a concentration of 150 μM LiTFSI. This expression was 1.4 times greater than the control group. Results revealed a decrease in the expression of *BAX* in HepG2 cells following exposure to LiTFSI. As the dosage increased, the expression of *BAX* showed a significant dose-dependent decrease.

Additionally, the expression of the four genes related to cell proliferation, Cyclin-dependent kinase inhibitor 1A (*CDKN1A)*, Cyclin-dependent kinase 4 (*CDK4)*, Ribosomal protein S6 (*RPS6), KIT* ligand *(KITLG)*, were analyzed. *CDKN1A* increased significantly at higher levels of LiTFSI exposure in both the cell lines, while *CDK4* exhibited a remarkable decrease. In A498 cells, there was no significant change in the expression of RPS6 after exposure to LiTFSI. Conversely, in HepG2 cells, a significant down-regulation of RPS6 was observed at high concentrations of LiTFSI. The expression of *KITLG* decreased at most concentrations in A498. *KITLG* expression decreased at 150 μM of LiTFSI treatment and increased at all higher concentrations in HepG2.

### 3.4 Dysregulation of Cell Cycle

Dysregulation of cell proliferation due to cell cycle abnormalities is a hallmark of tumorigenesis. To investigate whether LiTFSI influences cell proliferation by regulating cell cycle progression, we evaluated the expression of cell cycle genes in A498 and HepG2 cells (Figure 5a, Figure 5b, Figure S3 and Figure S4). Our results revealed that LiTFSI induced cell cycle arrest in the S phase in HepG2 cells, the proportion of cells in S phase increased by 67% at 150 μM LiTFSI compared to control. However, in A498 cells, LiTFSI did not affect cell cycle progression.

**Figure 5.**
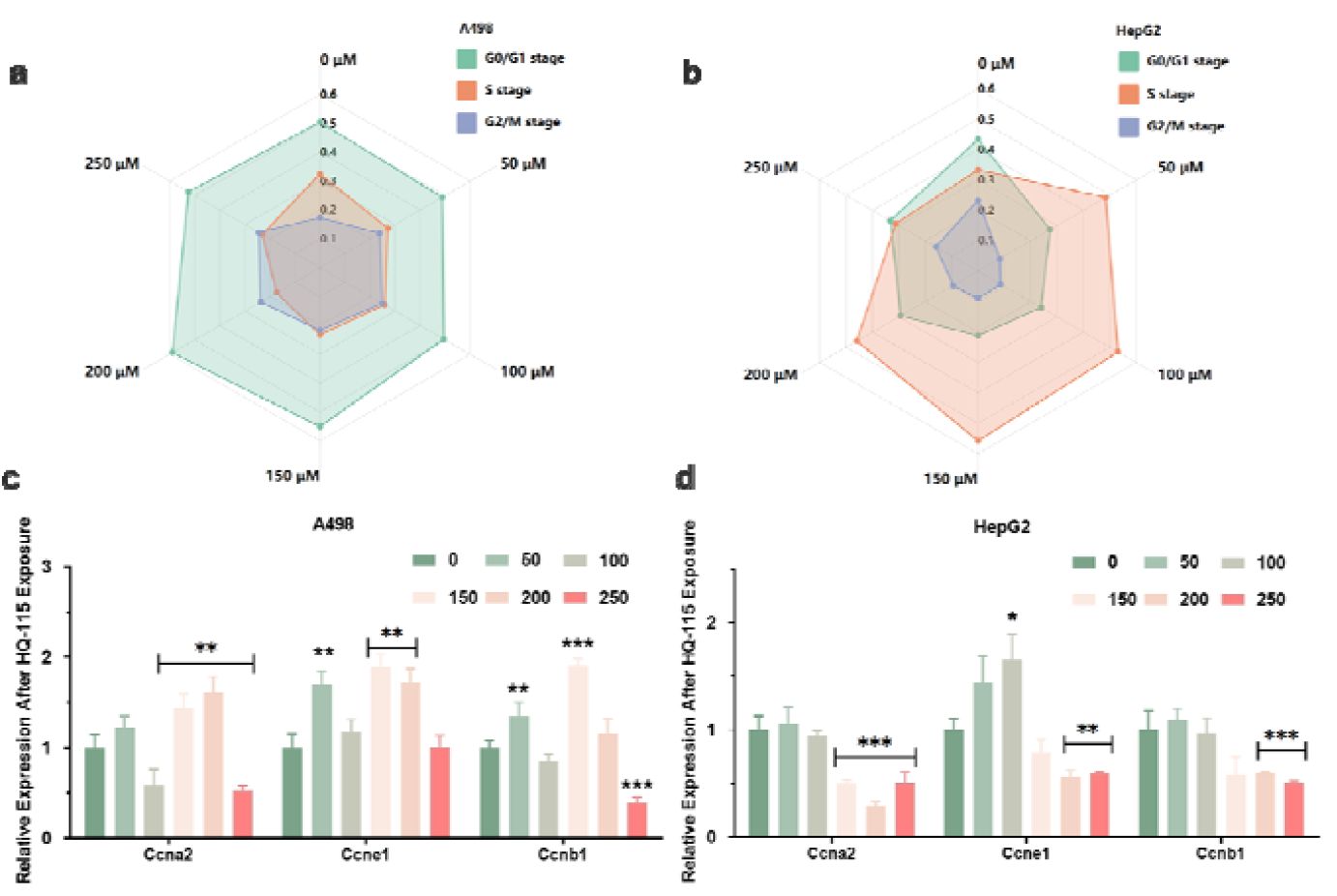
Cell cycle of A498 and HepG2 upon exposure to LiTFSI. (A) A498 (B) HepG2 cell cycle distributions detected by flow cytometry. Cells were exposed for 48 h in LiTFSI. Relative expression of cell cycle related genes after 48 h of exposure to LiTFSI (C) A498 (D) HepG2. Experiments were repeated three times. *p < 0.05 compared with untreated controls(NC).

The expression of cell cycle-related genes cyclin A2 (*Ccna2*), cyclin E1(*Ccne1*) and cyclin B1 (*Ccnb1*) were examined by qPCR to provide insights on the molecular mechanisms underlying the effect of LiTFSI on cell cycle progression. In our study, we observed differential regulation of *Ccna2, Ccne1*, and *Ccnb1* in A498 and HepG2 cells upon exposure to LiTFSI. In the A498 cells, the expression of *Ccna2, Ccne1*, and *Ccnb1* genes exhibited a significant upregulation at 150 μM of LiTFSI treatment, followed by downregulation at higher concentrations. In contrast, all three genes showed significant downregulation in HepG2 cells after treatment.

These findings suggest that the effects of LiTFSI on cell cycle progression may involve the selective modulation of specific cell cycle regulatory genes and is potentially cell type-dependent.

### 3.5 Effect on kidney damage and methylation genes

The carcinogenic potential of PFAS in humans has been reported in previous studies (Fenner, 2020; Pierozan et al., 2023; Shearer et al., 2021). Hence, the effect of PFAS on key genes associated with kidney injury, lipid metabolism and transportation as well as methylation was examined. A total of 19 mRNA expression levels were quantified and analyzed in this study.

The expression of genes associated with kidney injury and their response to LiTFSI exposure was investigated (Figure 6a, and Figure S5). Six genes were selected for evaluation and mRNA expression was performed on A498 cells. Among the genes studied, *Acta2* expression was significant decreased at LiTFSI doses of 100 μM and higher. *Tgfb1* and *Bcl2l1* exhibited a significant increase in gene expression only at a concentration of 150 μM, which was twice the control level, and insignificant changes were observed at other concentrations. In contrast, *Harvcr1, Nfe2l2*, and *Hes1* displayed significant increase expression at higher concentrations (>200 μM). The results provide evidence of a dose-dependent response of these genes to LiTFSI exposure and suggest that LiTFSI may have an impact on kidney function.

**Figure 6.**
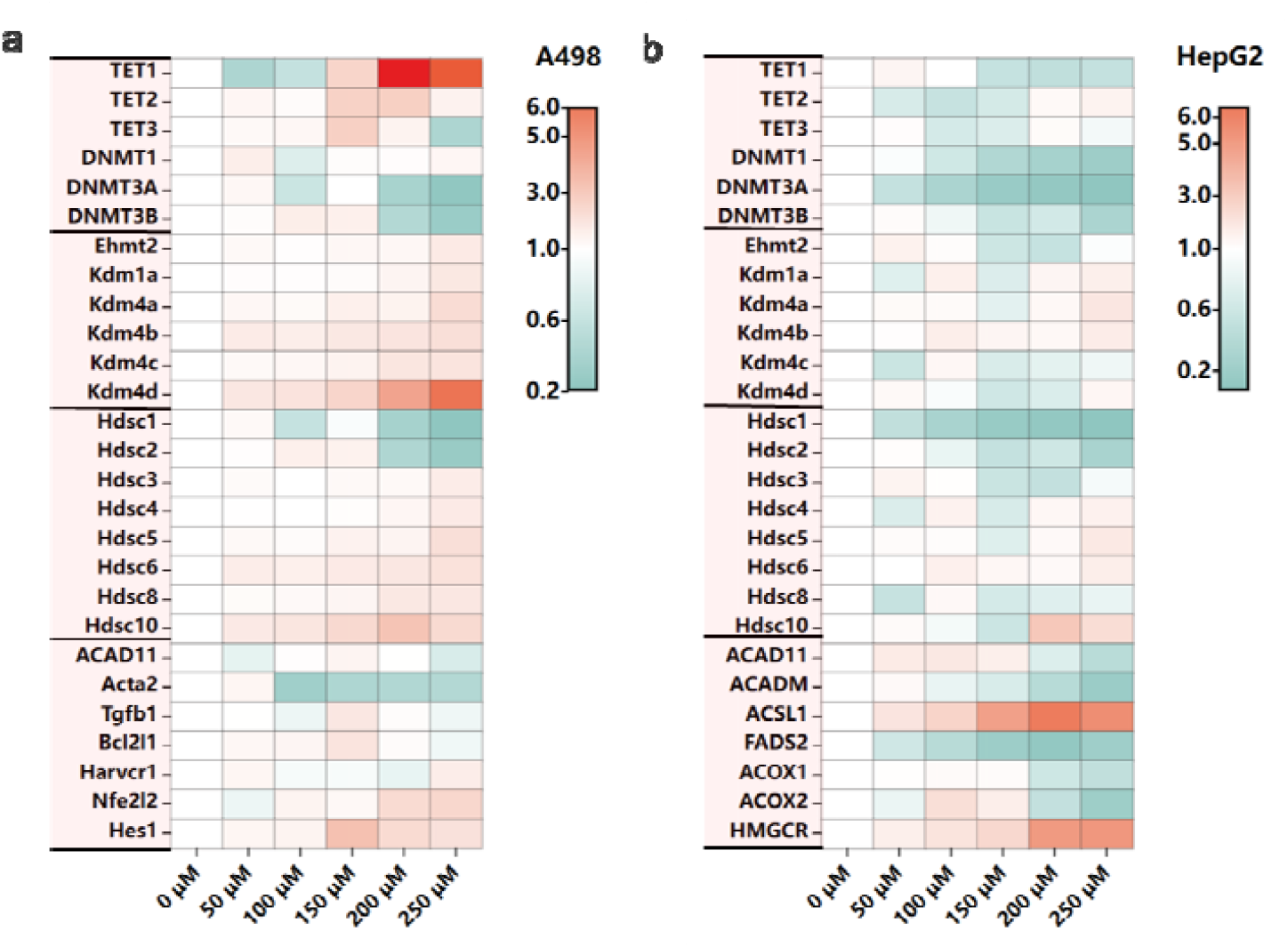
(A) Effect of LiTFSI on the expression of DNA methylation, histone methylation, histone acetylation and kidney damage related genes in A498. (B) Effect of LiTFSI on the expression of DNA methylation, histone methylation, histone acetylation and lipid metabolism, transportation related genes in HepG2. Experiments were repeated three times. *p < 0.05 compared with untreated controls(NC).

The methylation changes were investigated by assessing the relative gene expression profiles of *TET1, TET2, TET3, DNMT1, DNMT3A*, and *DNMT3B*, which are involved in the regulation of DNA methylation via *TET* methylcytosine dioxygenase and DNA methyltransferase enzymes. The expression of these genes was assessed in cells treated with different concentrations of LiTFSI (Figure S6). Our results revealed that with an increase in concentration of LiTFSI, the expression of *TET* family of genes exhibited an initial increase and subsequent decrease, with the highest expression observed at concentrations between 150 μM and 200 μM, which was more than 3 times higher than the control group. However, the expression of *DNMT1* was not significantly affected, while *DNMT3A* and *DNMT3B* were significantly down-regulated, both of which were less than half the control group. These findings suggest that LiTFSI may play a role in regulating the abundance of transcripts associated with DNA methylation through its effects on the expression of *TET* and *DNMT* genes.

### 3.6 Effect on lipid metabolism, transportation and methylation genes

Among the evaluated genes (Figure 6b), the expression of *ACAD11* and *ACADM*, which are involved in β-oxidation of fatty acids, showed a biphasic response upon exposure to LiTFSI, with an initial increase followed by a decrease. Conversely, the expression of *ACSL1*, a gene related to lipid synthesis significantly increased in a dose-dependent manner. *ACSL1* expression increased ∼ 6.5-fold upon exposure to 200 μM LiTFSI (Figure S7). *ACOX2*, encoding an enzyme involved in fatty acid degradation in peroxisomes, exhibited a significant reduction in its expression upon treatment with LiTFSI. Our results showed that exposure to LiTFSI at a concentration of 200 μM resulted in a significant increase in the expression of *HMGCR*, which encodes the rate-limiting enzyme in cholesterol synthesis. The fold change was approximately 5 times greater than the control group (Figure S7). Furthermore, we observed a clear modulation of transport genes related to lipid metabolism.

The expression of *TET* and *DNMT* family were assessed in cells exposed to different concentrations of LiTFSI (Figure S8). The expression of *TET1* and *TET3* displayed a downward trend (Less than 0.5 times) with increasing LiTFSI dose, while *TET2* expression was increased at higher concentrations of LiTFSI (250μM). Notably, the expression of all three *DNMT* genes showed a significant dose-response change.

The expression levels of the histone methyltransferase Ehmt2 and the demethylases Kdm1a, Kdm4a, Kdm4b, Kdm4c, and Kdm4d were assessed across all treatment groups. At all treatment levels, there was a significant increase in the expression of demethylases genes in A498 cells. However, for HepG2 cells, the increase in these genes was observed only at higher doses.

These results suggest that LiTFSI exposure could induce an increase in lipid anabolism and alterations in lipid catabolism in HepG2 cells. Our results shed light on the potential involvement of *TET* methylcytosine dioxygenases and DNA methyltransferases based on the observed methylation changes following exposure to LiTFSI in HepG2.

## 4. Discussion

Per- and polyfluoroalkyl substances (PFAS) have been noted to impart adverse effects on liver and kidney function (Evich et al., 2022; Wang et al., 2022). LiTFSI is designed for various industrial and commercial applications, including backup power systems, renewable energy storage systems, electric vehicles, and other high energy demanding applications (Ramesh and Ang, 2010). Despite the widespread use of LiTFSI in lithium-ion batteries and electronic devices, its toxicity has not been evaluated. Understanding the toxicity and persistence of LiTFSI will help inform future risk assessments and regulatory actions to protect human and environmental health. In the present study, we aim to elucidate the potential health risks of LiTFSI.

Compared with our previous study, LiTFSI was less cytotoxic than PFOA, PFOS, or GenX (Wen et al., 2020; Wen et al., 2022). The observed decrease in cell viability and increase in ROS production in both A498 and HepG2 cells following exposure to LiTFSI are consistent with previous studies which show that exposure to PFAS can induce oxidative stress and DNA damage in human cells (Dale et al., 2022; Ojo et al., 2021). However, the response of the two cell types to the oxidative stress is different, as indicated by their SOD expression levels. The upregulation of SOD expression in HepG2 cells indicates that these cells may mount an adaptive response to the oxidative stress induced by LiTFSI exposure. This increase in SOD expression allows the cells to effectively scavenge ROS and protect against oxidative damage which can lead to an increase in cell survival (Solan et al., 2023). However, the downregulation of SOD expression in A498 cells suggests that these cells may not be able to counteract the oxidative stress effectively. This decrease in SOD expression leads to an accumulation of ROS in the cells, which can cause oxidative damage and ultimately decrease cell viability. Therefore, the differential regulation of SOD expression in HepG2 and A498 cells suggests that HepG2 cells can better cope with oxidative stress upon exposure to LiTFSI, while A498 cells may be more susceptible to the negative effects of oxidative stress.

The induction of apoptosis in both cell lines is similar to previous studies which show that PFAS can induce programmable cell death (Bassler et al., 2019). Our findings also reveal that LiTFSI may modulate the expression of genes involved in the intrinsic and extrinsic apoptotic pathways in a cell type-specific manner. The study also evaluated the expression of genes related to apoptosis and proliferation in both cell lines. The expression of *BAX* (Bangma et al., 2020), a pro-apoptotic gene, was found to increase initially in A498 cells but decrease in a dose-dependent manner in HepG2 cells. In addition, the expression of *RPS6* is different in the two cell lines studied. *RPS6* may be involved in adipogenesis and promote the development of human hepatocellular carcinoma (Calvisi et al., 2011). The expression of *KITLG*, a gene involved in cell growth, decreased in A498 cells but increased in HepG2 cells. Taken together, our findings suggest that LiTFSI may modulate the expression of genes involved in apoptosis and proliferation. This could be due to differences in the expression of specific genes or proteins in the two cell types, or to differences in the way that LiTFSI interacts with these genes or proteins.

The modulation of cell cycle progression by LiTFSI is another important finding of this study. Previous research has indicated that PFOS, PFBS and PFBA could arrest cells in the G0 and G1 phases and decrease the number of cells in the S phase on bottlenose dolphin (Tursiops truncatus) skin cell (Otero-Sabio et al., 2022). The observation that LiTFSI induces cell cycle arrest in S phase in HepG2 cells but not in A498 cells suggests that the effects of LiTFSI on cell cycle progression are cell type-specific. The differential regulation of cell cycle-related genes *Ccna2, Ccne1*, and *Ccnb1* in A498 and HepG2 cells upon exposure to LiTFSI further supports this conclusion. This finding was also confirmed in HGrC1 cells (Clark et al., 2022).

Recent studies have shown that PFAS can impair lipid metabolism and cause damage to liver lipid homeostasis in frogs (Lin et al., 2022). Based on the experimental results, LiTFSI exposure caused significant changes in gene expression, indicating a potential impact on cellular function. The upregulation of genes involved in lipid metabolism, such as *ACSL1* and *HMGCR*, suggests a potential role in altering lipid synthesis and cholesterol levels (Zhao et al., 2022). The downregulation of genes involved in lipid catabolism, such as *ACOX2* and *ACAD11*, may indicate impaired fatty acid degradation (Kim et al., 2022). The changes in gene expression associated with kidney injury, such as upregulation of *Harvcr1* may promote cell death (Cheung et al., 2022). Moreover, *Nfe2l2* may increase the expression of antioxidant and detoxification genes, which could protect cells from damage caused by ROS (Jiang et al., 2022), consistent with the results of REDOX levels. The change in expression of *TET* and *DNMT* genes suggests that LiTFSI may influence the abundance of transcripts associated with DNA methylation (Wen et al., 2022), resulting in a significant downregulation of *DNMT3A* and *DNMT3B*. However, the effect of LiTFSI on the methylation level needs to be characterized in the future.

Overall, the results presented in this study demonstrate the short-term toxicity of LiTFSI in two different human cancer cell lines, A498 and HepG2. These findings suggest that LiTFSI, as a type of PFAS, can induce toxicity in human cells through a variety of mechanisms, including oxidative stress, apoptosis, and cell cycle dysregulation. These findings have important implications on the use of PFAS and related compounds in various applications, including con-sumer products and industrial processes. Future research directions could include investigating the long-term effects of LiTFSI exposure, studying the potential role of low-dose/chronic exposure to LiTFSI, and elucidating the molecular mechanisms underlying the cell type-specific effects of LiTFSI on various cellular processes.

## 5. Conclusions

Our work provides evidence on the short-term toxicity of LiTFSI, a novel PFAS. The results of the study showed that exposure to LiTFSI caused a decrease in cell viability and an increase in the production of ROS in both HepG2 and A498 cells. Our results are consistent with previous studies on in vitro toxicity of PFAS. Furthermore, LiTFSI was found to induce apoptosis in both the cell lines tested, and the expression of genes related to apoptosis and proliferation were affected upon exposure to LiTFSI. Additionally, LiTFSI was found to modulate cell cycle progression in a cell type-specific manner, which may have implications on therapeutic targets for cancer.

## Author Contributions

Conceptualization, J.I., J.G and X.Z.; methodology, X.Z.; software, X.Z. and M.S.; validation, M.L. and X.Z.; data curation, X.Z.; writing—original draft preparation, X.Z.; writing—review and editing, J.I., J.G. and X.Z.; supervision, J.I.; project administration, J.I. All authors have read and agreed to the published version of the manuscript.

## Institutional Review Board Statement

“Not applicable” Informed Consent Statement: “Not applicable.”

## Data Availability Statement

“Not applicable” Acknowledgments: Not applicable.

