## Supplementary Data File for "In vitro toxicity of LiTFSI on Human Renal and Hepatoma Cells"

**SUPPLEMENTARY MATERIALS**

**Contents list**

Additional Figure S1-S8 and Table S1

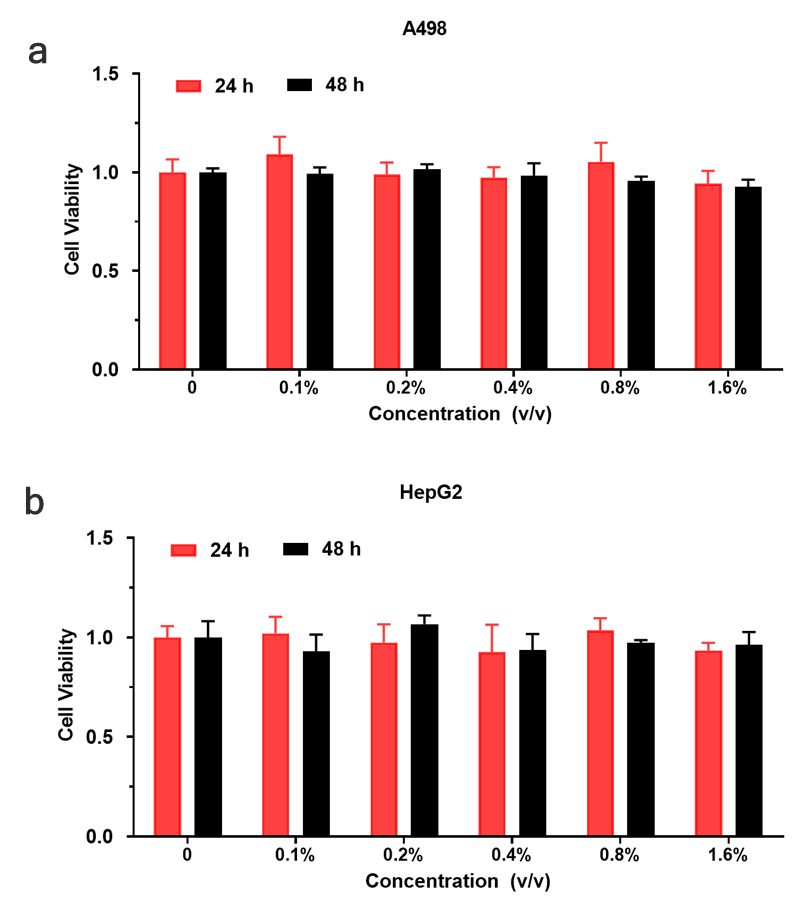

**Figure S1** Cytotoxicity assessment of DMSO in A498 and HepG2 cells. Experiments were repeated three times.

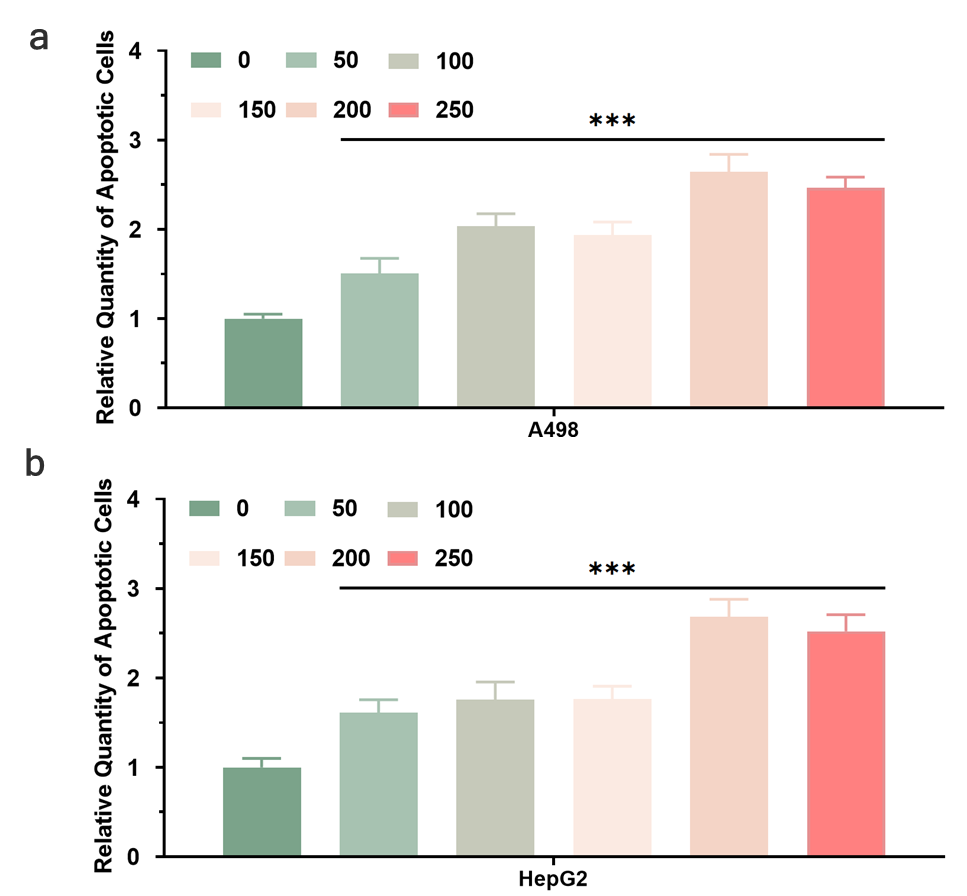

**Figure S2** Apoptosis evaluated by flow cytometry. Cells were exposed for 24 h in LiTFSI. Experiments were repeated three times. ∗p < 0.05 compared with untreated controls(NC).

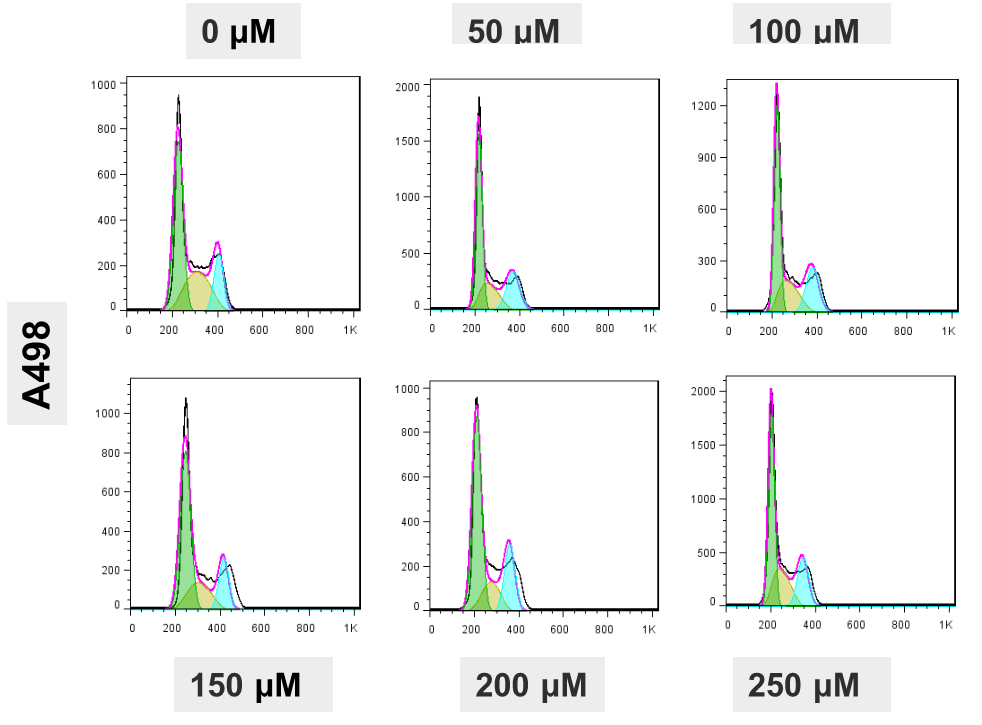

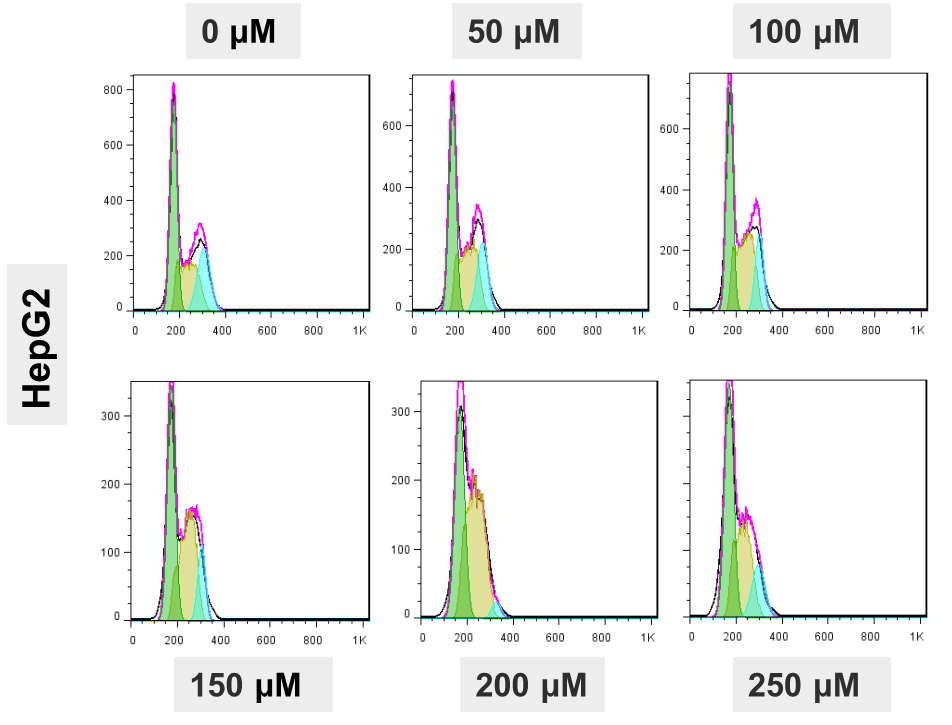

**Figure S3** Cell cycle evaluated by flow cytometry. Cells were exposed for 48 h in LiTFSI. Experiments were repeated three times.

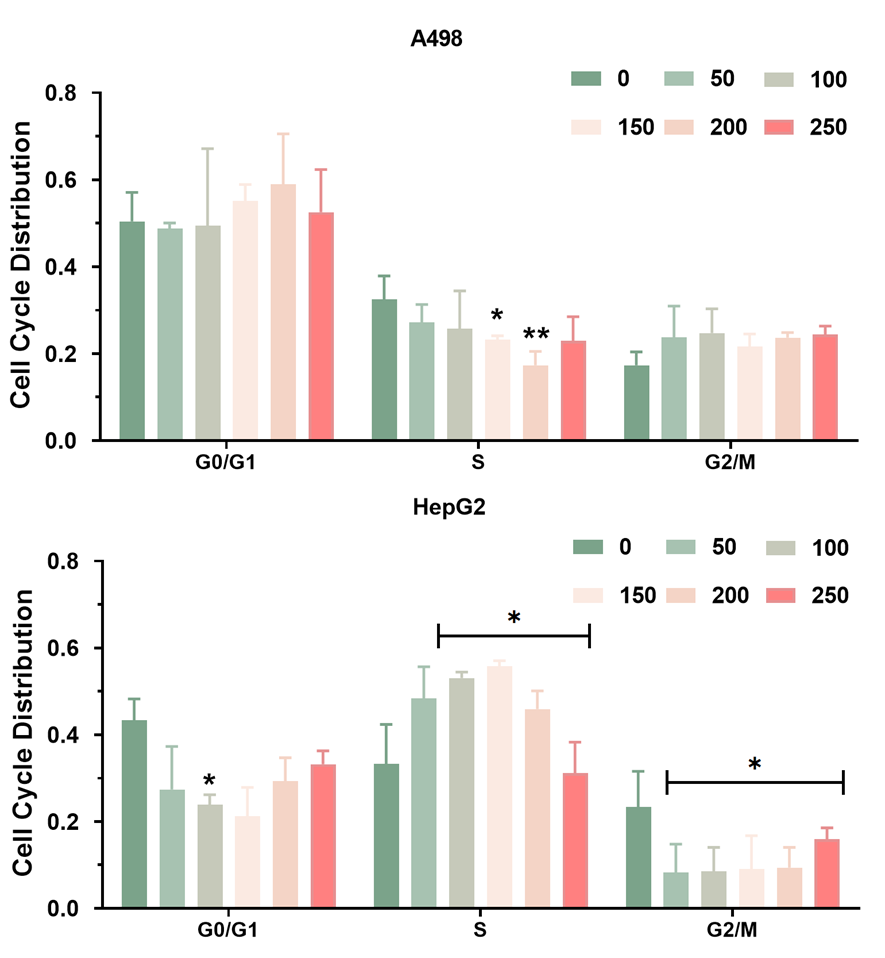

**Figure S4** Distribution of cell cycle. Cells were exposed for 48 h in LiTFSI. Experiments were repeated three times. ∗p < 0.05 compared with untreated controls(NC).

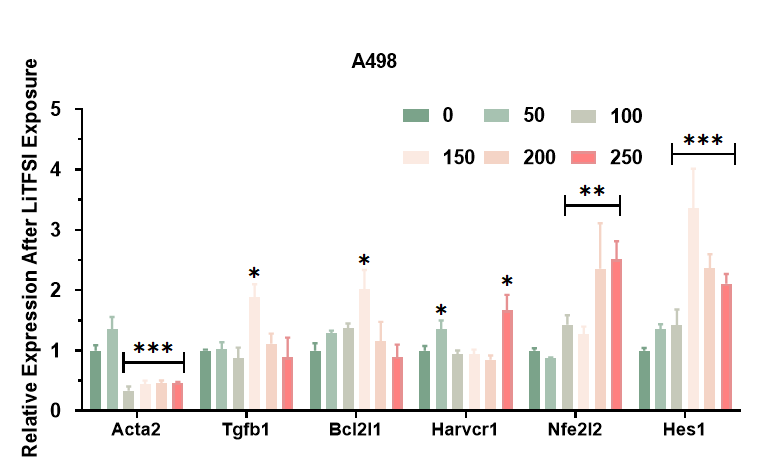

**Figure S5** Effect of exposure on the expression of kidney damage related genes in A498. Experiments were repeated three times. ∗p < 0.05 compared with untreated controls(NC).

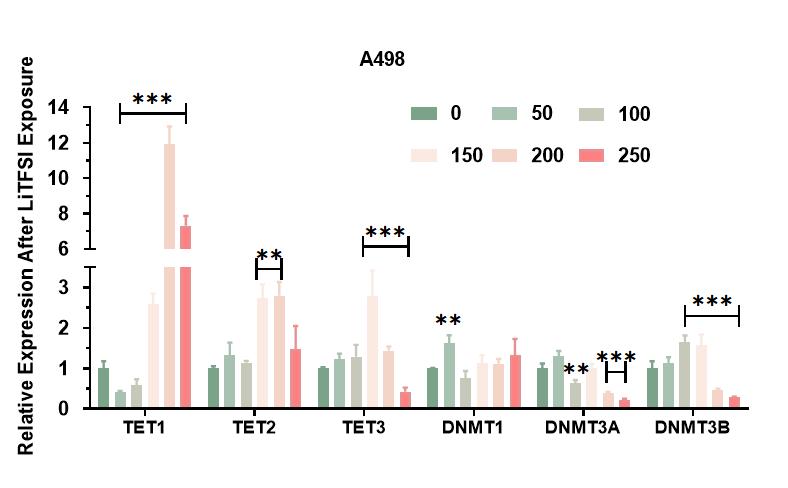

**Figure S6** Relative expression of Tet Methylcytosine Dioxygenases encoding TET1, TET2, and TET3, DNA Methyltransferases encoding DNMT1, DNMT3A, and DNMT3B after 24 h of exposure to LiTFSI in A498 cells.

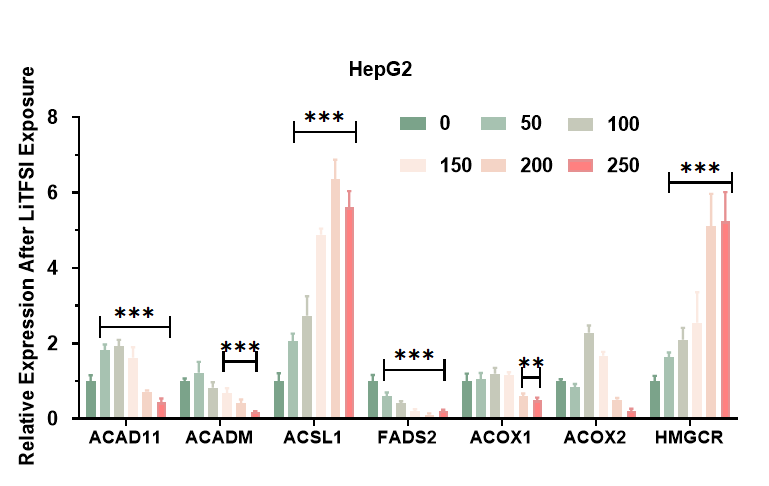

**Figure S7** Effect of exposure to LiTFSI on the expression of lipid metabolism and transportation related genes in HepG2 cells. Experiments were repeated three times. ∗p < 0.05 compared with untreated controls(NC).

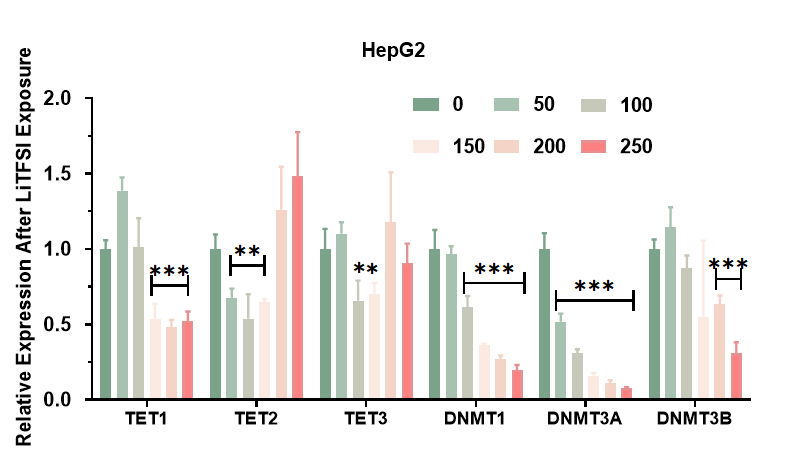

**Figure S8** Relative expression of Tet Methylcytosine Dioxygenases encoding TET1, TET2, and TET3, DNA Methyltransferases encoding DNMT1, DNMT3A, and DNMT3B after 24 h of exposure to LiTFSI in HepG2 cells.

**Table S1** Genes’ primer sequences designed for quantitative real-time polymerase chain reaction studies.

| Primer Name | Forward primer | Reverse primer |
| --- | --- | --- |
| *BAX* | CAAACTGGTGCTCAAGGCCC | GGGCGTCCCAAAGTAGGAGA |
| *CDKN1A* | TACCCTTGTGCCTCGCTCAG | GGCGGATTAGGGCTTCCTCT |
| *CDK4* | CTGTGCCACATCCCGAACTG | GCCTCTTAGAAACTGGCGCA |
| *RPS6* | CGATGAACGCAAACTTCGTA | TTCGGACCACATAACCCTTC |
| *KITLG* | GGCAAATCTTCCAAAAGACTACATG | CTACCATCTCGCTTATCCAACAATG |
| *CCNA2* | AGTAAACAGCCTGCGTTCACC | GAGGGACCAATGGTTTTCTGG |
| *CCNE1* | AAATGGCCAAAATCGACAGG | CGAGGCTTGCACGTTGAGTT |
| *CCNB1* | ATGACATGGTGCACTTTCCTCC | GCCAGGTGCTGCATAACTGG |
| *ACAD11* | CAGGGTCGAATCTTCCGTGAT | GTCCTGATGAGCTGCAGCTT |
| *ACADM* | GAGCAGGCTCTGATGTAGCTG | TTCCTGGGGTATCTGCTTCC |
| *ACSL1* | GACCTCTCCATGCAGTCAGT | GACACCTGTATTCCCCTCTGG |
| *FADS2* | CTGACCTGGAATTCGTGGGC | CATGTCCTCAGCCGTCTTCC |
| *ACOX1* | CAGCCTGAAAGCTGAGTTGC | TCTGGGTGCAGGAAGAAGAA |
| *ACOX2* | CTTTCTGGAGACGTGGCCTT | GGCCCTGAAGATATGTCCCATG |
| HMGCR | TGGGAATGCAGAGAAAGGTGC | AAGCTCCCATCACCAAGGAG |
| Acta2 | GGAGGAGATGACGAGCCCTT | GTTCCAGCTTTCCAGGGAGT |
| Tgfb1 | GGGACACCCACTGCAAGAC | CAGCTCCACGCAGGATTT |
| Bcl2l1 | GCTGGGGCCAACTCTAAGAT | CACAGGGTCATTTCCACTTG |
| Harvcr1 | CGCGCGCCTGCTCGT | CCATCCAGCGCTGAATGTTT |
| Nfe2l2 | GACAGCGACTGAGGAGGAGA | CGCGAGGTTCTCTGACTGTT |
| Hes1 | TACCTGGAGAGCCGAGTTTG | TGTGTTTCCGCTTCCAGGTA |
| GAPDH | GAAGGTGAAGGTCGGAGTC | GAAGATGGTGATGGGATTTC |
